# Combined celecoxib and NDGA induces a non-canonical, autophagy-independent redox cell death via p62/KEAP1 in colorectal cancer

**DOI:** 10.64898/2026.09.29.753356

**Authors:** Rajesh Yadala, Sharvani Palakurthy, Smita C. Pawar

## Abstract

Colorectal cancer (CRC) exhibits substantial intratumoral heterogeneity, with differences in oncogenic driver mutations and apoptotic competence contributing to therapeutic resistance. Here, we investigated whether the combined celecoxib (a COX-2 inhibitor) and NDGA (a 5- LOX inhibitor) treatment (C+N) induces redox-mediated cell death via the p62/KEAP1 axis, offering a therapeutic approach capable of transcending genetic diversity in CRC. We used two genetically distinct CRC cell lines, HCT116 and HT29, harbouring KRAS and BRAF mutations respectively, with differing p53 status, as models of tumor heterogeneity. C+N exhibited profound cytotoxicity in both cell lines, independent of COX-2/5-LOX expression and apoptotic competence. Additionally, C+N induced marked oxidative stress with mitochondrial ROS, lipid peroxidation, and labile ferrous iron accumulation, visualized as distinct JC-1 green mitochondrial puncta and FerroOrange red iron puncta in both cell lines. C+N-induced death was reversed by the antioxidant NAC but not by inhibitors of apoptosis, autophagy, necroptosis, or ferroptosis, indicating a mechanism distinct from these pathways. Mechanistically, combination induced p62 Ser349 phosphorylation, promoting KEAP1 degradation and NRF2 stabilization independently of ULK1-mediated autophagy; however, NRF2 stabilization was uncoupled from HO-1 induction. Our work reveals that C+N induces a ULK1/autophagy-independent, redox-mediated cell death mechanism that circumvents apoptotic competence, highlighting redox disruption as a genotype-independent therapeutic strategy to overcome CRC heterogeneity.

## Introduction

Colorectal cancer (CRC) is the second leading cause of cancer-related death worldwide and the third most frequently diagnosed malignancy, with 1.93 million new cases and 903,859 deaths recorded in 2022 [1, 2]. Although advanced screening and multimodal therapy have shown betterment in localized disease, the prognosis of metastatic CRC remains poor, mainly due to therapeutic resistance developed by use of chemotherapy and targeted therapy [3].

A major factor contributing to treatment failure is intratumoral genetic heterogeneity [4, 5]. This heterogeneity is markedly seen in KRAS (40–50%) and BRAF (∼10%) mutant CRC, which exhibit increased metabolic reprogramming [6–8]. Furthermore, another factor contributing to differential drug sensitivity is the mutational status of p53 on apoptotic responses [9–11]. To capture this heterogeneity, we selected HCT116 (p53 wild-type, KRAS-mutant) and HT29 (p53-mutant, BRAF-mutant) cells, allowing us to assess whether any potent therapeutic strategy remains effective independent of apoptotic competence.

Cancer cells exhibit abnormal metabolism, mitochondrial dysfunction, and oncogenic signaling and, thereby, need to be facilitated with robust antioxidant defenses. This dependence on antioxidant capacity is a vulnerability, where excessive oxidative stress can overwhelm cellular defenses and can ultimately lead to cell death [13, 14]. In recent years, the altered redox metabolism of cancer cells has emerged as an attractive target to develop therapeutic strategies. Oxidative stress can engage multiple regulated cell death modalities beyond apoptosis, autophagy-dependent death, necroptosis, and ferroptosis, and the iron-dependent form of death driven by lipid peroxidation [15]. Given this diversity of possible outcomes, therapeutic strategies that induce oxidative stress may act through mechanisms that extend beyond, or circumvent, apoptosis altogether.

The KEAP1-NRF2 pathway is the master regulator of cellular redox homeostasis. Under basal conditions, KEAP1 promotes the ubiquitin-dependent proteasomal degradation of NRF2; however, under oxidative stress, this interaction is disrupted, leading to the nuclear translocation of NRF2 and transcriptional activation of cytoprotective genes [16–17]. NRF2 activation through the non-canonical pathway is mediated by phosphorylation of p62 at Serine 349 that further enhances KEAP1 sequestration, resulting in NRF2 release and stabilization [18, 19]. Given that severe oxidative insults can activate the non-canonical p62/KEAP1/NRF2 pathway, we investigated whether this pathway was involved in the context of drug-induced oxidative stress.

Celecoxib, a selective cyclooxygenase-2 (COX-2) inhibitor, exerts anticancer effects in CRC through both COX2-dependent [20] and independent [21] mechanisms, including mitochondrial superoxide production, membrane depolarization, and ROS-dependent apoptosis [21, 22]. Despite chemopreventive efficacy in familial adenomatous polyposis [23], celecoxib has not significantly improved overall survival in CRC patients. COX-2 inhibition can shunt arachidonic acid metabolism toward the 5-lipoxygenase (5-LOX) pathway [24]. Nordihydroguaiaretic acid (NDGA), a natural lignan and 5-LOX inhibitor, similarly exerts antineoplastic effects through modulation of oxidative stress and inflammatory pathways, as well as through mechanisms independent of LOX inhibition [25]. Based on this rationale, we hypothesized that combined celecoxib and NDGA treatment would enhance cytotoxicity in CRC cells.

Here, we show for the first time that combined treatment with celecoxib and NDGA (C+N) induces redox-mediated cell death via the non-canonical p62/KEAP1 axis, independent of COX-2/5-LOX in CRC cell lines.

## Methods

### Cell lines

Human CRC cell lines HCT116 (KRAS mutant, p53 wild-type) and HT-29 (BRAF and p53 mutant) were obtained from the National Centre for Cell Sciences (Pune, India). HCT116 cells were cultured in McCoy’s 5A media (Cytiva, USA), and HT29 cells were cultured in DMEM (Cytiva, USA) supplemented with 10% FBS (Gibco, Thermo Fisher Scientific) in a humidified incubator at 37°C with 5% CO_2._

### Reagents and antibodies

Celecoxib (T0466), NDGA (TJS2190), Licofelone (T6568), Z-VAD-FMK (T7020),

Necrostatin-1 (T1847), 3-MA (T1879), and NAC (T0875) were purchased from TargetMol (Boston, MA, USA). RSL3 (HY-100218A) and Ferrostatin-1 (HY-100579) were purchased from MedChem-Express (Monmouth Junction, NJ, USA). Primary antibodies against PARP (detecting full-length and cleaved forms) and HO-1 were purchased from CST (Danvers, Massachusetts, USA). NRF-2 (A3577), KEAP-1 (A25951), SQSTM1/p62 (A19700), Phospho-SQSTM1/p62-S349 (A20737), and ALOX5 (A2877) phosphor-ULK1 S555 (AP1495) were purchased from ABclonal (Woburn, MA, USA). COX-2 (sc-376861) antibody was purchased from Santa Cruz (Finnell Street, Dallas, Texas, USA). RIPA buffer (R0278), phosphatase inhibitor (P0044), and protease inhibitor (P8340) were purchased from Sigma-Aldrich, Burlington, Massachusetts, USA.

### CCK-8 cell viability assay

To determine cell viability, 3000-5000 cells per well were seeded in a 96-well plate. After treatment with the compounds for 24 h, 10 µl of CCK-8 (TargetMol, Boston, MA, USA) was added to each well and incubated in a humidified incubator at 37°C with 5% CO_2_ for 3 h. Absorbance was measured at 450 nm wavelength.

### Colony formation assays

Colony formation assay was performed to assess clonogenic survival. Cells were seeded at low density in 6-well plates and cultured for 7–14 days following treatment. The resulting colonies were fixed with ice-cold 100% methanol for 15 minutes at room temperature and stained with 0.5% (w/v) crystal violet for 30 minutes. Plates were washed with running tap water and air-dried.

### Annexin V-FITC/7-AAD apoptosis assay

Cells were incubated with 5 µl of Annexin V-FITC and 5 µl of 7-AAD (E-CK-A212, Texas, USA) for 20 mins and were run in a Beckman Cytoflex flow cytometer. The results were analyzed using Cytoflex software v.2.0.

### DCFH-DA cellular ROS assay

Dichlorofluorescein diacetate (DCFH-DA) assay was performed to measure ROS by flow cytometer. Cells were incubated for 10 mins with 10 µM DCFH-DA in Hank’s balanced salt solution and were analysed using a Beckman CytoFlex flow cytometer.

### Measuring Mitochondrial Superoxide Using MitoSOX™ Green

Cells were incubated with MitoSOX™ Green (M36006, Thermo Fisher Scientific) for 30 mins at a concentration of 1 µM in Hank’s balanced salt solution in a humidified incubator at 37°C with 5% CO_2_. Cells were analyzed using a Beckman CytoFlex flow cytometer, and data were plotted using FlowJo software (version 10).

### Assessment of Mitochondrial Membrane Potential Using TMRM

Mitochondrial membrane potential was assessed using the fluorescent cationic dye Tetramethylrhodamine Methyl Ester (TMRM) (17109, TargetMol, Boston, MA, USA). Cells were incubated with 50 nM TMRM in culture medium for 30 minutes at 37°C. Following incubation, cells were washed, resuspended in PBS, and analyzed immediately by a CytoFLEX flow cytometer (Beckman Coulter, USA).

### Lipid ROS (reactive oxygen species) assay by C11-BODIPY® 581/591

Cells were incubated with C11-BODIPY® 581/591 (D3861, Thermo Fisher Scientific USA) for 30 mins at a concentration of 2 µM in Hank’s balanced salt solution in a humidified incubator at 37°C with 5% CO_2_. Cells were analyzed using a Beckman CytoFlex flow cytometer, and data were analysed using FlowJo software (version 10).

### Intracellular iron measurement by FerroOrange

Intracellular iron levels were quantified using FerroOrange (F374-10, Dojindo Laboratories, Japan), a specific fluorescent probe for labile Fe (II). Cells were incubated in the dark with 2 µM FerroOrange in Hanks’ Balanced Salt Solution (HBSS) at 37°C for 30 min. Cellular fluorescence (as an indicator of Fe (II) content) was visualized using an EVOS 7000 fluorescence microscope (Thermo Fisher Scientific, USA).

### Protein isolation and western blotting

Total protein was isolated by lysing cells on ice for 30 min in RIPA buffer containing protease and phosphatase inhibitors and vortexed intermittently. The lysates were centrifuged at 12,000 rpm for 30 min at 4 °C and protein concentrations were determined by BCA assay. Equal amounts of protein (30 μg) were separated by SDS–PAGE and blotted onto nitrocellulose membranes. Membranes were blocked with 5% skimmed milk for 1 h at room temperature and incubated overnight at 4 °C with primary antibodies, followed by 1 h incubation at room temperature with HRP-conjugated secondary antibodies. After washing with TBST, the protein signals were detected with ECL (G-Biosciences, St. Louis, USA) and imaged with a ChemiDoc system (Bio-Rad, California, USA).

### Statistical analysis

Statistical analysis was performed using GraphPad Prism 8.0 software (San Diego, CA, USA). All data are mentioned as mean ± standard deviation (SD). Comparisons between two groups were analyzed using unpaired Student’s t test, while multiple group comparisons were assessed by one way ANOVA followed by Sidak’s multiple comparisons test. A p value of < 0.05 was considered statistically significant, with *p < 0.05, **p < 0.005, ***p < 0.001, and ****p < 0.0001.

## Results

### Celecoxib and NDGA combination exhibit COX-2/5-LOX-independent cytotoxicity in CRC

Two CRC cell lines were exposed to different concentrations of celecoxib, NDGA, licofelone, and RSL3. Licofelone (a dual COX-2/5-LOX inhibitor) and RSL3 (a ferroptosis inducer) served as positive controls in our study. Based on dose-response assays (Fig. S1), IC_50_ values were determined (Table S1). For subsequent experiments, HCT116 and HT29 cells were treated with celecoxib (40 µM), NDGA (70 µM), and their combination. The C+N combination induced greater cell death than either agent alone in both cell lines (Fig. 1A).

**Fig. 1.**
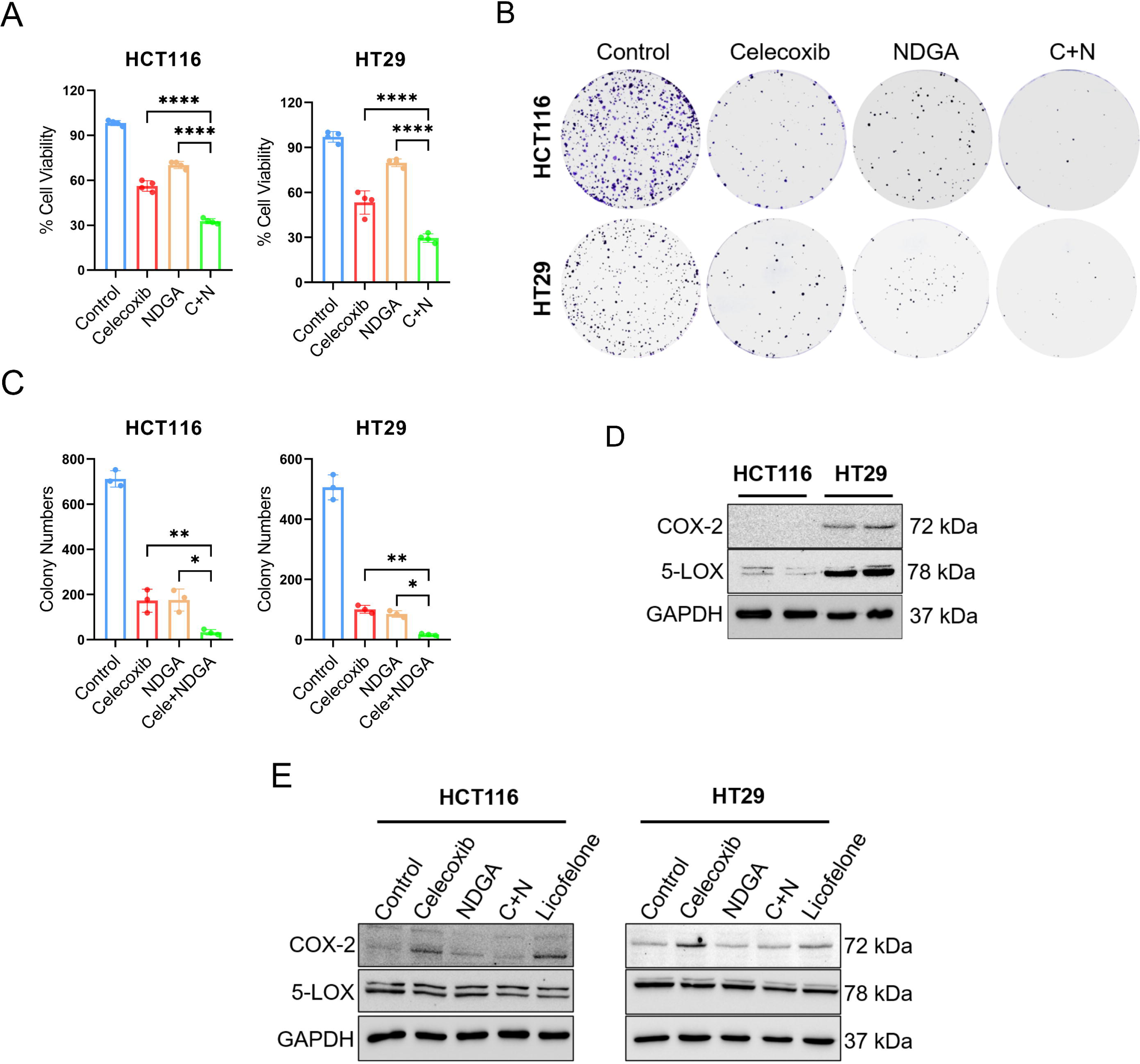
Celecoxib and NDGA Combination exhibit COX-2/5-LOX-independent cytotoxicity in CRC. **A** HCT116 and HT29 cells were treated with celecoxib (40 µM), NDGA (70 µM), or their combination (C+N) for 24 hours. Cell viability was assessed by CCK-8 assay and expressed as a percentage of the untreated control (n=4 each). **B** Colony formation assay with the indicated treatments was performed (left panel), and a histogram of colony numbers is shown (right panel). Data are shown as mean ± SD (n=3) (\**p* < 0.02; \*\**p* < 0.005 versus celecoxib alone or NDGA alone). **C** Western blot analysis of basal COX-2 and 5-LOX protein levels in HCT116 and HT29 cells. **D** Western blot analysis of COX-2 and 5-LOX protein levels in HCT116 and HT29 cells after treatment with the indicated compounds.

Consistent with these cytotoxicity findings, C+N significantly suppressed clonogenic survival in both CRC cell lines compared to individual treatments (Fig. 1B-C). To further investigate the antiproliferative effects, we evaluated the dose-dependent effects of celecoxib, NDGA, and their combination on clonogenicity at various concentrations. C+N showed a profound decrease in colony survival at all given concentrations compared to individual drugs in both cell lines (Fig. S2).

We next examined the basal protein levels of COX-2 and 5-LOX in both CRC cell lines by western blot analysis. HCT116 cells exhibited undetectable or very low expression of both proteins, whereas HT29 cells showed comparatively high basal expression (Fig. 1D). Treatment with C+N did not alter COX-2 or 5 LOX levels in either cell line. Surprisingly, celecoxib or licofelone increased COX-2 expression in both CRC cells, while NDGA did not induce any change (Fig. 1E). HCT116 cells, which lack detectable expression of COX-2 and 5LOX proteins, remained sensitive to C+N treatment. Collectively, these findings confirm that the cytotoxic effects of C+N occur independently of COX 2 and 5 LOX expression.

### NDGA shifts celecoxib-induced cell death away from classical apoptosis in HCT116 cells

Annexin V-FITC/7-AAD staining revealed that celecoxib and licofelone significantly increased the early apoptotic population (Annexin V**^+^**/7-AAD□) in HCT116 cells; however, NDGA alone did not significantly increase the Annexin V signal, and notably, the C+N failed to enhance and instead diminished the apoptotic signal compared to celecoxib treatment alone. In contrast, HT29 cells exhibited no Annexin V-positive signal across all treatment groups (Fig. 2A).

**Fig. 2.**
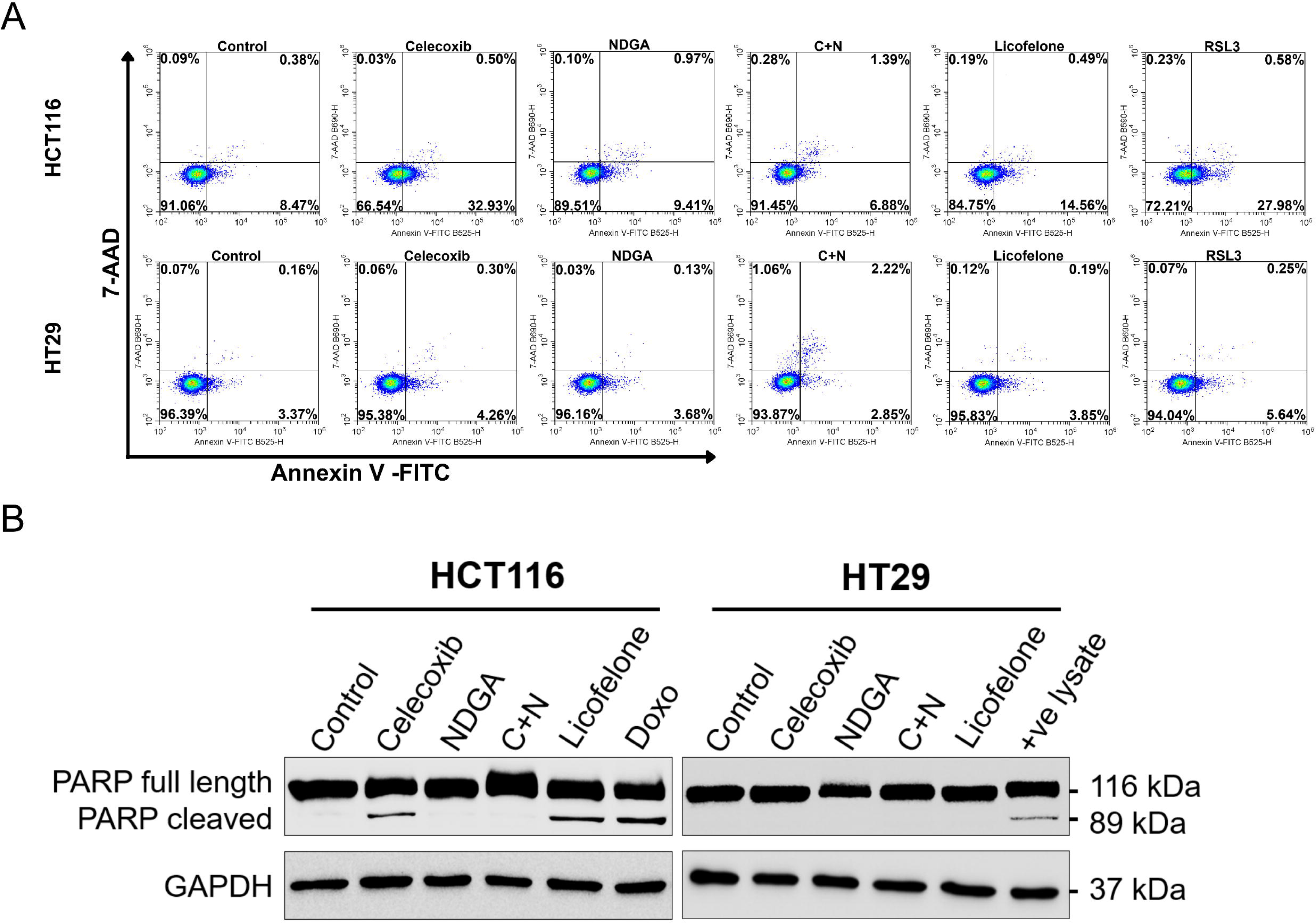
NDGA shifts celecoxib-induced cell death away from classical apoptosis in HCT116 cells. **A.** HCT116 and HT29 cells were treated with the indicated compounds for 24 hours. Apoptosis was assessed by Annexin-V FITC/7-AAD staining and analyzed by flow cytometry. The percentage of apoptotic cells is shown. **B** Western blot analysis of cleaved PARP levels in HCT116 and HT29 cells after treatment with the indicated compounds. Doxorubicin-treated HCT116 cell lysate was used as a positive control for cleaved PARP (last well in the HT29 panel).

To confirm these observations, cleaved PARP, a marker of apoptosis, was assessed. Consistent with Annexin V data, cleaved PARP was detected only in HCT116 cells treated with celecoxib or licofelone. Co-treatment with NDGA (C+N) abolished celecoxib-induced PARP cleavage, indicating that NDGA antagonizes rather than potentiating celecoxib-induced apoptotic signaling (Fig. 2B). Given that C+N retains its cytotoxic efficacy despite this loss of apoptotic marker (Fig. 1A-B), these results suggest that NDGA shifts C+N-induced cell death mechanism away from classical apoptosis. No cleaved PARP was detected in HT29 cells under any treatment conditions, confirming the absence of a robust apoptotic response in this p53 mutant cell line (Fig. 2B).

### C+N triggers redox imbalance and lipid peroxidation

To further charcaterize the metabolic changes induced by C+N, we first assessed intracellular ROS levels using DCFH-DA. In both cell lines, C+N induced a significant increase in DCFH-DA fluorescence compared to either agent alone. Additionally, licofelone induced ROS in both cell lines (Fig. 3A).

**Fig. 3.**
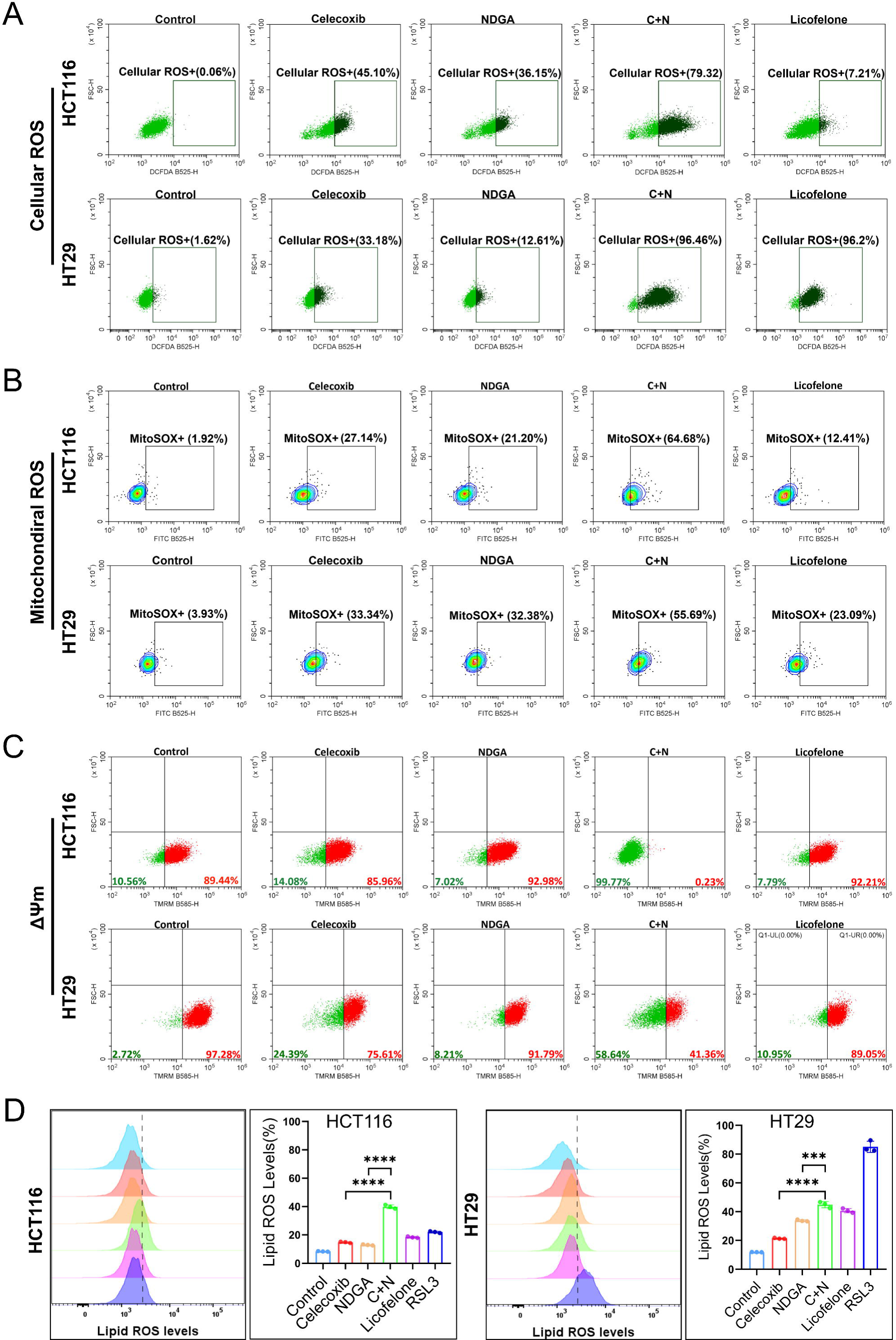
C+N triggers redox imbalance and lipid peroxidation. **A** Representative flow cytometry plots showing global ROS levels measured by DCFDA staining in HCT116 and HT29 cells treated with the indicated compounds for 24 hours. The percentage of ROS positive cells is indicated within each plot. **B** Representative flow cytometry plots showing mitochondrial superoxide (O□•□) production measured by MitoSOX™ Green staining in HCT116 and HT29 cells treated with the indicated compounds for 24 hours. The percentage of MitoSOX positive cells is indicated within each plot. **C** Representative flow cytometry plots showing mitochondrial membrane potential (ΔΨm) measured by TMRM staining in HCT116 and HT29 cells treated with the indicated compounds for 24 hours. Red indicates cells with intact membrane potential, while green indicates cells with loss of ΔΨm. The percentage of cells in each quadrant is indicated within the plots. **D** Lipid peroxidation was measured using C11 BODIPY® 581/591 staining followed by flow cytometry in HCT116 and HT29 cells treated with the indicated compounds for 24 hours. Data are shown as mean ± SD (n=3). \*\*\**p* = 0.0001; \*\*\*\**p* < 0.0001 versus single agents (celecoxib or NDGA alone).

Given the central role of mitochondria in redox signalling, we quantified the mitochondrial superoxide (O□•□) using MitoSOX™ Green and mitochondrial membrane potential (ΔΨm) using TMRM. C+N produced a pronounced increase in MitoSOX fluorescence accompanied by profound loss of ΔΨm, indicating potent mitochondrial superoxide generation and membrane depolarization. Licofelone increased MitoSOX in both cell lines but caused only a modest decrease in ΔΨm in HT29 cells, with no effect in HCT116 cells (Fig. 3B-C).

To additionally evaluate the oxidative response, we quantified lipid peroxidation using C11- BODIPY 581/591. C+N significantly elevated lipid ROS levels relative to either agent alone (Fig. 3D) in both cell lines. RSL3 and licofelone also increased in lipid ROS levels in both cell lines.

### Redox-dependence of C+N-mediated cell death: rescue by NAC but insensitive to Z-VAD-FMK, 3-MA, Necrostatin-1, and Ferrostatin-1

To determine the type of cell death induced by C+N, we performed pharmacological rescue experiments using cell death-specific inhibitors. Cells were pretreated for 2 hours and subsequently co-treated for 24 hours with the respective treatments in the presence of Z-VAD-FMK (pan-caspase inhibitor), 3-methyladenine (3-MA, autophagy inhibitor), Necrostatin-1 (necroptosis inhibitor), Ferrostatin-1 (ferroptosis inhibitor), and N-acetylcysteine (NAC, antioxidant). Evident with the substantial increase in lipid ROS levels in C+N (Fig. 3D), we included RSL3 as a positive control.

In HCT116 cells, Z-VAD-FMK significantly rescued cells in celecoxib and licofelone treatment, confirming the apoptotic nature in this p53 wild-type cell line (Fig. 4A), while in HT29 cells, Z-VAD-FMK failed to rescue any treatment group. Neither 3-MA, necrostatin-1, nor ferrostatin-1 rescued cells in the given drug treatments in either cell line, with the exception of RSL3 induced death, which was rescued by ferrostatin-1 as expected (Fig. 4B- D). Together, these results exclude apoptosis, autophagy, necroptosis, and ferroptosis as the primary cell death mechanism underlying C+N-induced cell death. In sharp contrast, NAC provided substantial protection across all treatment groups in both cell lines, confirming that oxidative stress is a central and unifying driver of cell death (Fig. 4E). Together, these findings establish that C+N induces a redox-dependent form of cell death that is mechanistically distinct from well-characterised regulated cell death pathways.

**Fig. 4.**
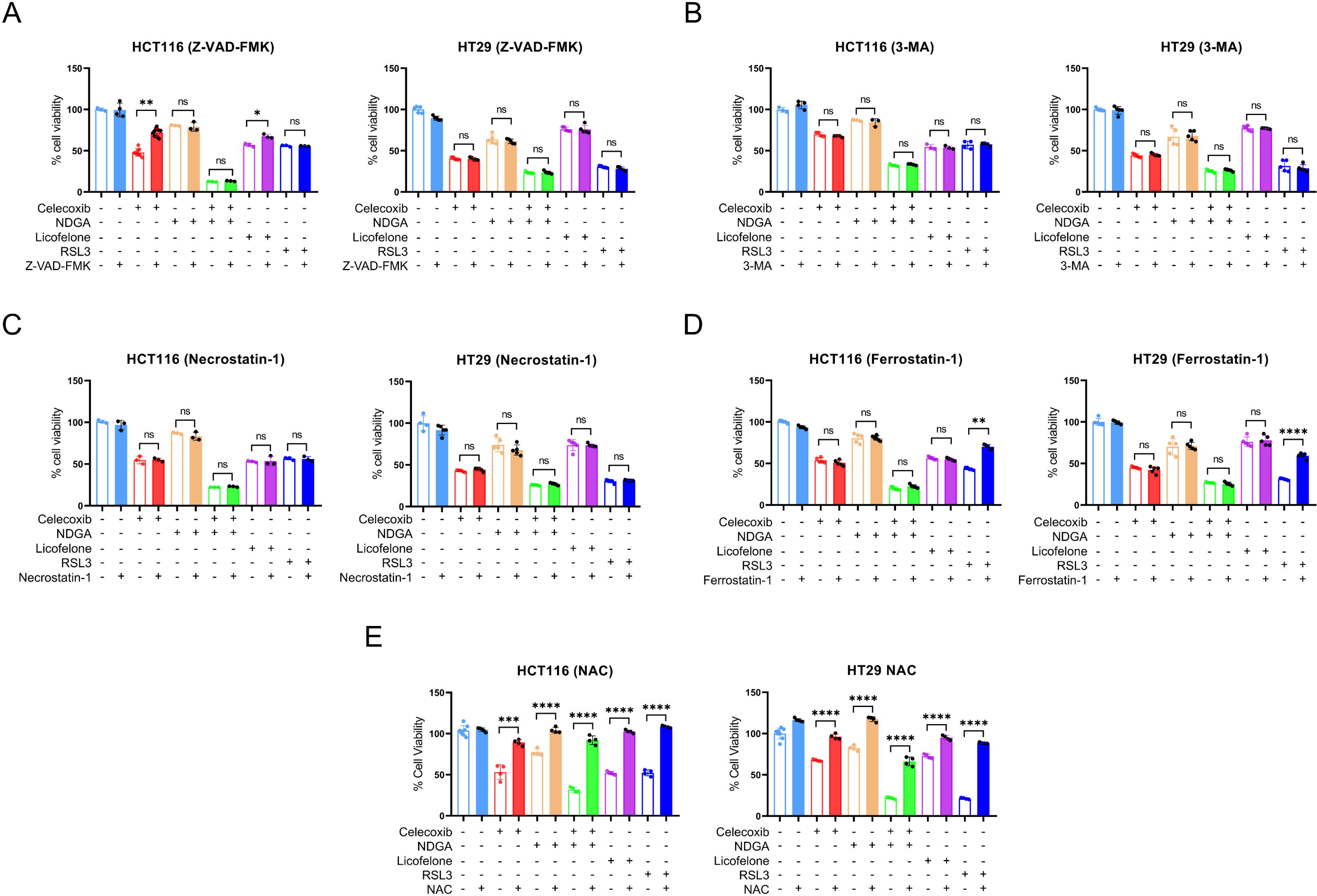
Redox-dependence of C+N-mediated cell death: rescue by NAC and insensitivity to Z-VAD-FMK, 3-MA, Necrostatin-1, and Ferrostatin-1. HCT116 and HT29 cells were pretreated with the indicated inhibitors for 1 hour and subsequently co-treated with the respective compounds for 24 hours. Cell viability was assessed by CCK-8 assay. A Z-VAD- FMK (10 µM) B 3-MA (60 µM) C Necrostatin-1 (10 µM) D Ferrostatin-1 (10 µM) (ferroptosis inhibitor), and E N-acetylcysteine (5 mM) (NAC, antioxidant). Data are indicated as mean ± SD (n=3). \**p* < 0.05; \*\**p* = 0.005; \*\*\**p* = 0.0001; \*\*\*\**p* < 0.0001.

### C+N induces green mitochondrial puncta and red iron puncta, reflecting a redox-dependent cellular phenotype

To further characterize this redox-driven phenotype at the subcellular level, we assessed intracellular iron homeostasis and mitochondrial integrity by fluorescence imaging.

Ferrorange staining revealed that C+N-treated cells exhibited a unique feature of localized red Ferrorange puncta. The bright red granular structures indicate a very high concentration of labile ferrous iron (Fe^2+^) within discrete cytosolic compartments and thus are a visual demonstration of intracellular redox dysregulation caused by this drug combination (Fig. 5A).

**Fig. 5.**
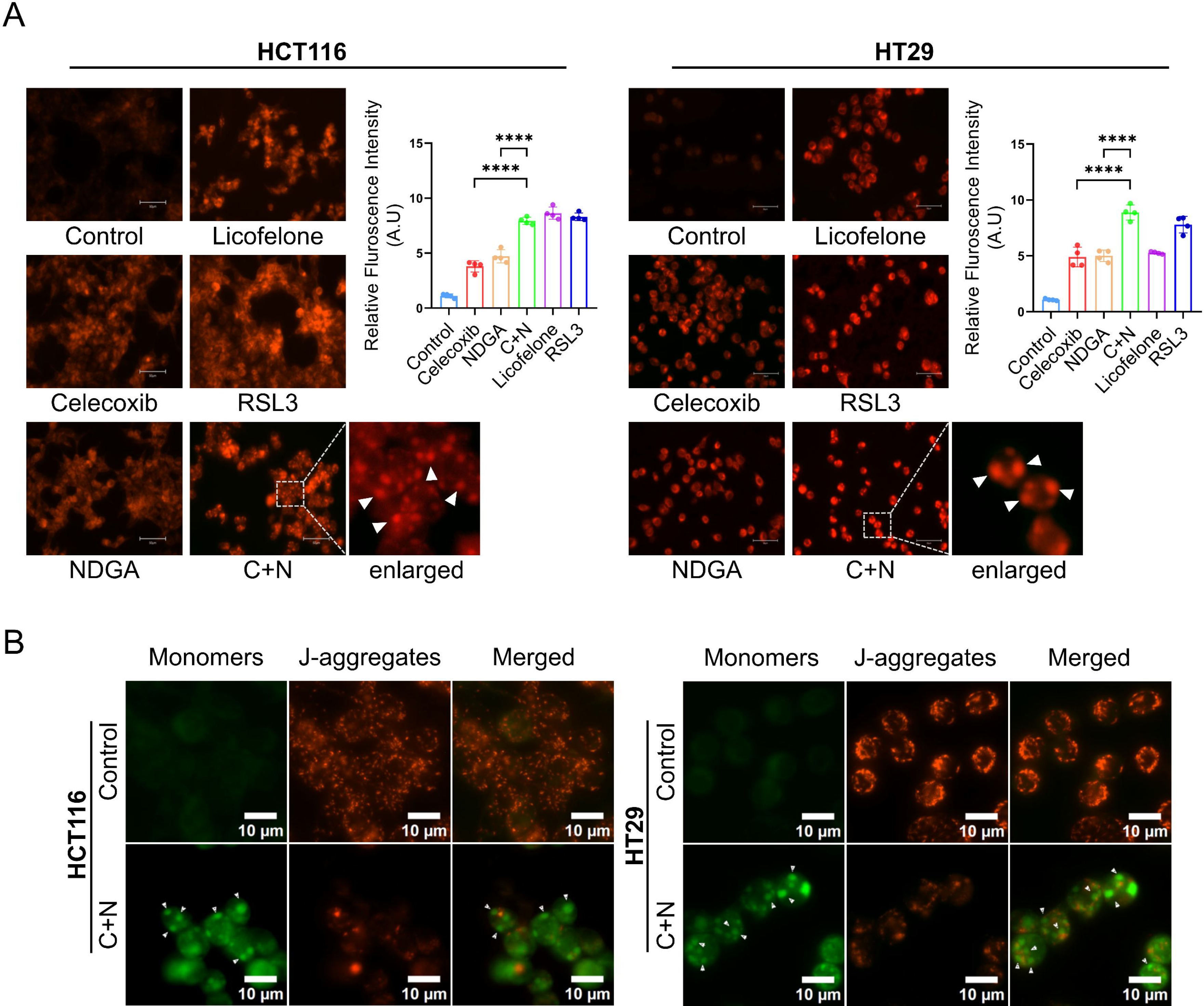
Redox-dependent cellular phenotype characterized by green mitochondrial puncta and red iron puncta. **A** Representative fluorescence microscopy images showing mitochondrial morphology by JC-1 staining. Red fluorescence represents healthy, hyperpolarized mitochondria, and green fluorescence indicates depolarized mitochondria. Arrows specify green mitochondrial puncta. **B** Representative fluorescence microscopy images showing intracellular labile Fe (II) levels estimated by FerroOrange staining. Red fluorescence indicates accumulation of labile ferrous iron. Arrows specify red iron puncta. Scale bar, 10 μm. Data are shown as mean ± SD (n=4). \*\*\*\**p* < 0.0001 versus single agents (celecoxib or NDGA alone).

Similarly, in untreated cells, JC-1 exhibited a robust red fluorescent signal, indicative of functional, hyperpolarized mitochondrial networks. However, the C+N combination caused a striking shift in the morphology of the mitochondria wherein its reticular networks underwent profound fragmentation, characterized by the formation of discrete, bright green JC-1 monomeric puncta (Fig. 5B). This abrupt transition from a red polarized network to green depolarized puncta represents a direct subcellular outcome of severe mitochondrial oxidative damage.

These green mitochondrial puncta and red iron puncta were consistently formed in the HCT116 and HT29 cell lines after C+N treatment (Fig. 5A-B). These morphological alterations define a prominent redox-associated phenotype within the cell, characterized by concomitant intracellular iron accumulation and mitochondrial fragmentation.

### C+N induces redox-mediated cell death via p62-KEAP-1 axis

p62 (also known as sequestosome 1, SQSTM1) is a multifunctional stress-inducible protein that regulates oxidative stress response through the KEAP1–NRF2 pathway [26]. Established that C+N induces oxidative stress-mediated cell death; we further examined the p62/KEAP1 axis. Western blot analysis revealed a marked increase in phosphorylation of p62 at Ser349 following C+N treatment in both cell lines, while total p62 levels remained relatively unchanged (Fig. 6A, 6B).

**Fig. 6.**
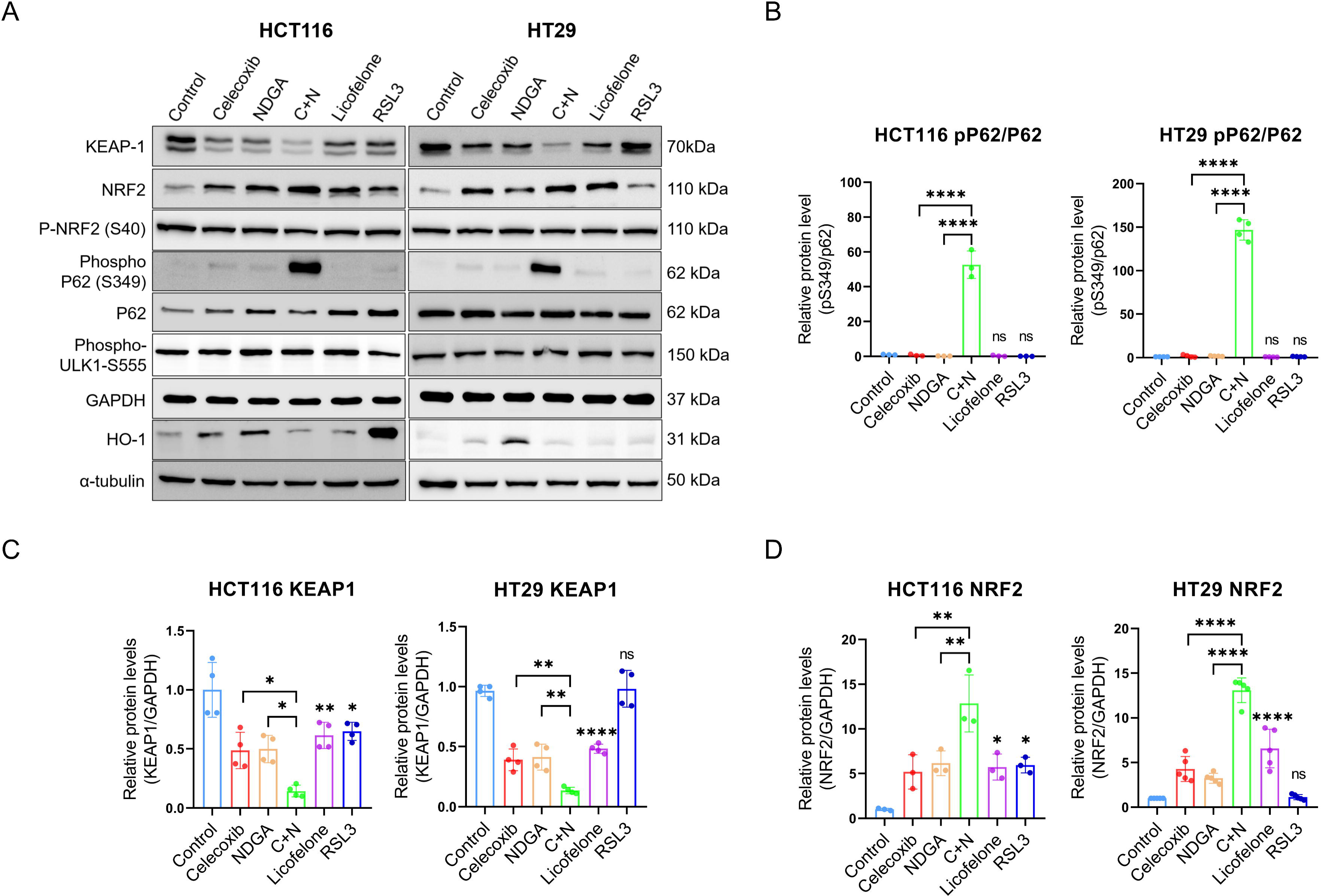
Celecoxib/NDGA combination induces redox-mediated cell death via non canonical phosphorylation of p62 (S349) followed by activation of NRF2 and degradation of KEAP-1. A Western blot analysis of total p62, phospho-p62 (S349), KEAP1, NRF2, and phospho-ULK1 in HCT116 and HT29 cells after 24 hours of treatment with the mentioned compounds. GAPDH served as a loading control. B-D Densitometric quantification of phospho-p62/total p62 ratio, KEAP1, and NRF2 in HCT116 and HT29 cells. Values were normalized to GAPDH. Data are shown as mean ± SD (n=3). \**p* < 0.05; \*\**p* < 0.005; \*\*\**p* < 0.0005; \*\*\*\**p* < 0.0001 versus single agents (celecoxib or NDGA alone).

Since phosphorylation of p62 at Ser349 increases its affinity for KEAP1, promoting KEAP1 sequestration, degradation, and subsequent NRF2 stabilization [27], we next assessed KEAP1 levels. KEAP1 was decreased by celecoxib, NDGA, licofelone, and C+N in both cell lines, with RSL3 producing a similar effect in HCT116 but not in HT29 cells; C+N produced a significant decreased compared to either agent alone (Fig. 6A, 6C). Since KEAP1 degradation is expected to stabilize NRF2, we next examined NRF2 and its activating phosphorylation Ser40 [27]. NRF2 was increased by the same treatments that decreased KEAP1, with the greatest increase again observed in C+N (Fig. 6A, 6D); however, phosphorylation at Ser40 remained unchanged across all treatment groups (Fig. 6A), indicating that this increase in total NRF2 was not accompanied by its activating phosphorylation. We next examined whether NRF2 activated its target gene, heme oxygenase-1 (HO-1) [28]. HO-1 induction was cell line and treatment specific. In HCT116 cells, celecoxib, NDGA, and RSL3 increased HO-1, whereas C+N and licofelone did not; in HT29 cells, only NDGA induced HO-1 (Fig. 6A). Together, these results show that although C+N increases total NRF2, this is not accompanied by Ser40 phosphorylation or HO-1 induction.

Given p62’s role in selective autophagy [19], we hypothesised that the strong p62 phosphorylation by C+N is associated with the activation of autophagic signaling. However, phosphorylation of ULK1 at Ser555, an activating modification for autophagy induction [29], was not significantly increased after C+N treatment (Fig. 6A), suggesting that p62-mediated KEAP1 sequestration is unlikely to be mediated by ULK1-dependent autophagy.

Although cells appear to mount a p62/KEAP1-dependent antioxidant response by profound phosphorylation of p62, the strong oxidative insult caused by C+N and the absence of significant NRF2/HO1 imply that this response may be functionally insufficient to overcome the oxidative burden caused by cell death observed after combination treatment.

Collectively, our results indicate that the C+N combination activates the p62/KEAP1 axis via noncanonical p62 phosphorylation and KEAP1 degradation in a redox-dependent and ULK1- independent manner.

## Discussion

CRC remains a leading cause of cancer-related death. Conventional therapeutic strategies targeting DNA synthesis or oncogenic drivers are often nonspecific and limited by resistance. The elevated redox state of cancer cells presents an alternative therapeutic vulnerability, since drugs that exacerbate oxidative stress can selectively eliminate malignant cells across genetically distinct tumors [12]. Our study identifies that combined celecoxib and NDGA (C+N) treatment exerts antiproliferative effects on CRC cells in a redox-dependent manner, through activation of a noncanonical p62/KEAP1 axis that fails to resolve into productive antioxidant signaling.

Celecoxib and NDGA have individually been reported to exert anticancer effects through mechanisms beyond their canonical targets, including Wnt/β-catenin signaling, PDK1/AKT signaling, GSK3-dependent c-FLIP degradation, mitochondrial superoxide production, and ROS-dependent apoptosis for celecoxib [30–33], and TRAF6–IL-6–STAT3, IGF-1R/HER2, GSK-3β/NRF2 signaling, and glutathione depletion for NDGA [34–37]. To our knowledge, however, combined celecoxib and NDGA treatment has not previously been reported in CRC in a COX-2/5-LOX-independent context. Prior work showed that COX-2 inhibition shunts arachidonic acid metabolism toward 5-LOX, and that combined COX-2/5-LOX inhibition (celecoxib plus AA-861) suppresses tumor growth more effectively than either agent alone [38]. In contrast, we found that celecoxib and licofelone increased COX-2 expression in both cell lines, while NDGA and C+N left COX-2 and 5-LOX levels unchanged, consistent with reports that NSAIDs can upregulate COX-2 via PPAR-mediated mechanisms [39]. This induction, however, does not necessarily reflect increased enzymatic activity, as HT29 cells express catalytically inactive COX-2 with minimal PGE output [40]. Together with the persistence of C+N cytotoxicity in HCT116 cells, which lack detectable COX-2/5-LOX expression, these findings confirm that C+N acts independently of these canonical targets.

p53-mutant cells are resistant to apoptosis per se, limiting the effectiveness of conventional chemotherapeutics such as 5-FU and oxaliplatin [41–42]. Consistent with this, HT29 cells (p53-mutant) showed no cleaved PARP under any treatment, whereas HCT116 cells (p53 wild-type) did (Fig. 2B), indicating that classical apoptotic responses are genotype-restricted in this system. Importantly, C+N produced comparable cytotoxicity in both cell lines, despite this difference, suggesting that its mechanism circumvents apoptotic pathways entirely, a feature that may be particularly advantageous in apoptosis-resistant colorectal tumors.

The pronounced increase in lipid ROS and labile Fe² accumulation observed with C+N is a hallmark feature typically associated with ferroptosis and initially suggested this iron-dependent death pathway as the likely driver of cytotoxicity. However, the failure of ferrostatin-1 to rescue C+N-treated cells asserts against this interpretation, indicating that lipid peroxidation and iron accumulation in our model arise as downstream consequences of redox dysregulation rather than as drivers of ferroptosis. Combined with the inability of Z-VAD-FMK, 3-MA, and necrostatin-1 to give protection and the complete rescue of cells by NAC, these findings suggest that C+N-induced death cannot be attributed to any single canonical regulated cell death pathway and is instead governed fundamentally by redox imbalance itself.

The p62-KEAP1-NRF2 axis a well-characterized cytoprotective pathway in which Ser349 phosphorylation of p62 displaces NRF2 from KEAP1 and activates antioxidant genes including HO-1 [19, 43]. Ikeda et al. recently showed that ULK1 phosphorylates p62 at Ser349 within p62 bodies, trapping KEAP1 for autophagic degradation and thereby stabilizing NRF2 to drive HO-1 expression and cell survival [44]. Our findings diverge from this model in two key respects. First, phosphorylation of ULK1 at Ser555 was unchanged by C+N, indicating that ULK1 is not activated in our system. Second, the autophagy inhibitor 3- MA failed to rescue C+N-induced death, confirming that autophagy is not engaged.

However, we did not observe any HO-1 induction despite this NRF2 stabilization, suggesting that NRF2 is transcriptionally uncoupled from its canonical target, in direct contrast to the Ikeda et al. model in which NRF2 activation leads to robust HO-1 expression and survival [44]. Furthermore, phosphorylation of NRF2 at Ser40 also remained unchanged, consistent with Relic et al. [45] who reported that the expression of constitutive p-NRF2 (S40) does not increase upon NRF2 stabilization. This failure to mount a productive antioxidant response probably created a permissive environment for the unchecked oxidative stress we observed: elevated cytosolic ROS, mitochondrial superoxide, lipid peroxidation, and loss of mitochondrial membrane potential. We therefore propose that C+N triggers a ULK1/autophagy-independent redox-driven cell death pathway mediated by KEAP1 depletion and stabilization of NRF2, but its transcriptional uncoupling from HO-1 prevents an effective antioxidant response, permitting catastrophic ROS accumulation and cell death. This converts the p62-KEAP1-NRF2 axis from a survival mechanism into a mediator of redox-driven death, representing a cancer-specific vulnerability that may be therapeutically exploitable, consistent with emerging evidence that targeting tumor antioxidant systems can induce lethal oxidative stress in cancer cells [46].

This mitochondrial and redox dysfunction was accompanied by a distinct subcellular phenotype as C+N treatment induced accumulation of labile ferrous iron alongside JC-1 green puncta indicative of mitochondrial fragmentation and depolarization, consistently across both cell lines. Iron metabolism is frequently dysregulated in CRC, with malignant cells accumulating higher iron levels to support cellular activity while requiring tight regulation of redox-active iron to prevent oxidative damage [47–48]. Ponnusamy et al. recently demonstrated that CRC cells possess an adaptive iron-buffering mechanism that restricts expansion of the labile iron pool, even when iron synergizes with cell death inducers, pointing to intracellular iron compartmentalization as a protective mechanism [49]. The increased intracellular iron accumulation observed during combination treatment may be an adaptive response to sequester or compartmentalize iron and limit iron-mediated oxidative damage by C+N.

The therapeutic exploitation of oxidative stress is being explored across multiple cancers. Arsenic trioxide is FDA-approved for leukemia [50], isoalantolactone increases CRC cell sensitivity via ROS and mitochondrial dysfunction [51], and vitamin C selectively targets KRAS/BRAF-mutant CRC cells through excessive ROS generation [52]. Other redox-targeting agents, including ATN-224, 2-methoxyestradiol, curcumin derivatives, carnosol, and 15d-PGJ2, are similarly under investigation as pro-oxidant anticancer strategies [53–57]. In this context, C+N induced significant loss of mitochondrial membrane potential and ROS generation in mitochondria suggesting its potential as a redox-targeting strategy in CRC with the advantage of functioning independently of cell’s apoptotic sensitivity.

To summarize, our results demonstrate the therapeutic potential of combined celecoxib and NDGA treatment in heterogeneous tumors resistant to apoptosis-based and targeted therapies. Given the established clinical safety profiles of celecoxib and NDGA, the anticancer potential of C+N through redox modulation presents a unique strategy for CRC therapy. Further mechanistic and in vivo studies are needed to better understand this cell death pathway.

## CRediT authorship contribution statement

**Smita C. Pawar:** Conceptualization, Funding Acquisition, Supervision, Validation, Writing - Original Draft, Writing - Review & Editing. **Rajesh Yadala:** Conceptualization, Data Curation, Formal Analysis, Investigation, Methodology, Project Administration, Validation, Visualization, Writing - Original Draft, Writing - Review & Editing. **Sharvani Palakurthy:** Investigation, Methodology, Writing - Original Draft, Writing - Review & Editing.

## Acknowledgments

The authors thank the Department of Genetics and Biotechnology, Osmania University, for providing infrastructure support. **Rajesh Yadala** acknowledges Indian Council of Medical Research for providing Senior Research Fellowship (ICMR-SRF) [File no: 45/24/2020/PHA/BMS]. **Smita Pawar** acknowledges the support from Department of Biotechnology (DBT)-BUILDER Programme [grant number BT/INF/22/SP41415/2021]; the Department of Science and Technology (DST) FIST Level II [grant number SR/FST/LS- II/2022/985]; and the Anusandhan National Research Foundation (ANRF) [grant number EEQ/2023/001015].

## Data availability

Data will be made available on request.

## Graphical Abstract

The graphical abstract was created using https://BioRender.com.

## Declaration of Generative AI and AI-assisted technologies in the writing process

ChatGPT (OpenAI) was used only to refine grammar and improve the readability of the written text. All scientific content, analysis, and conclusions were independently developed and verified by the authors, who take full responsibility for the published work.

## Conflict of Interest

The authors declare that they have no known competing financial interests or personal relationships that could have appeared to influence the work reported in this paper.

---

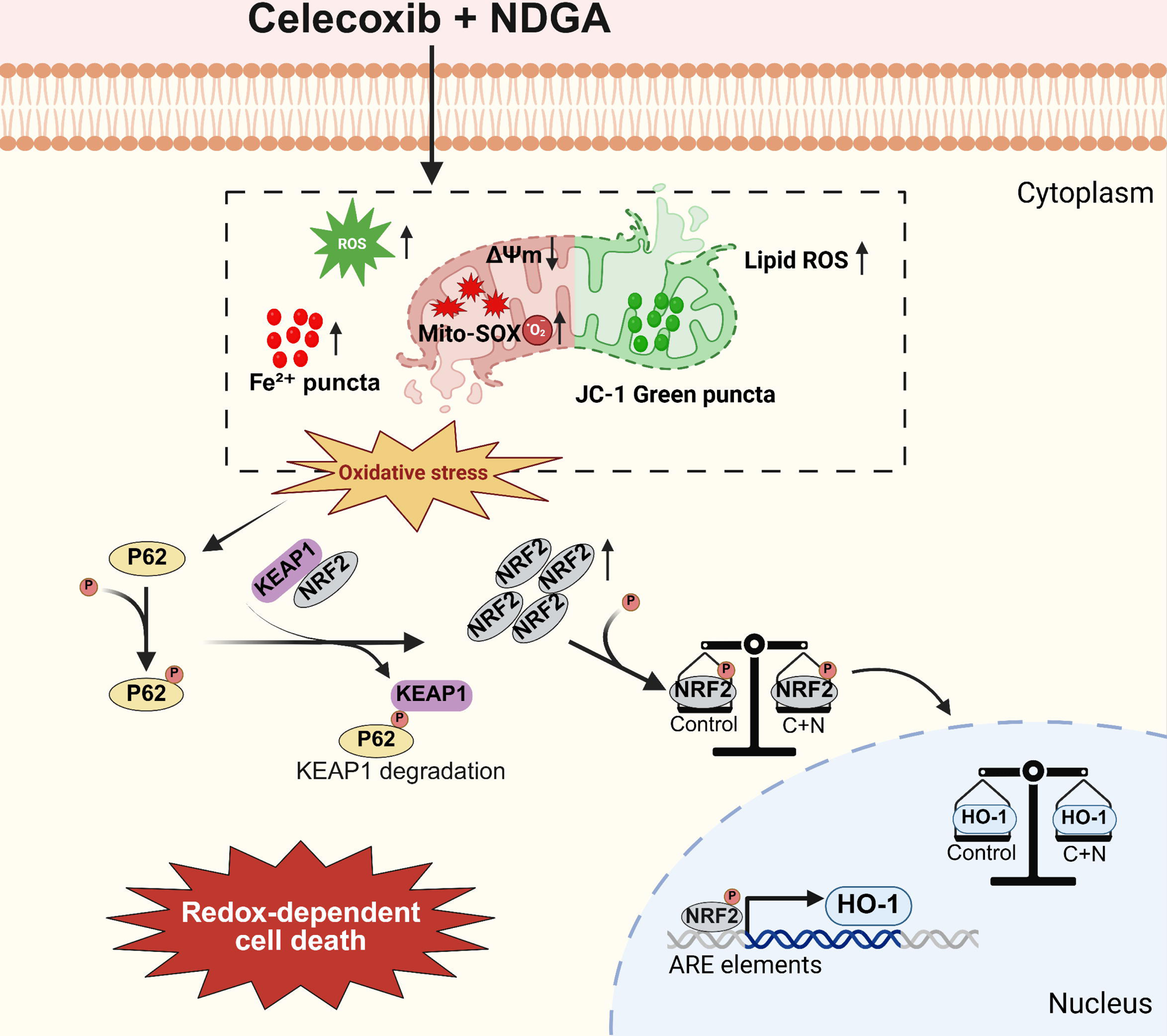

## Notes

### Competing Interest Statement

The authors have declared no competing interest.

### Summary of Updates

The second author's name was changed from Sharvani palakuthi to Sharvani Palakurthy

