## Supplementary figures and images for "Combined celecoxib and NDGA induces a non-canonical, autophagy-independent redox cell death via p62/KEAP1 in colorectal cancer"

### Supplemental data 1

#### Uncropped Western Blots:

Fig. 1 D

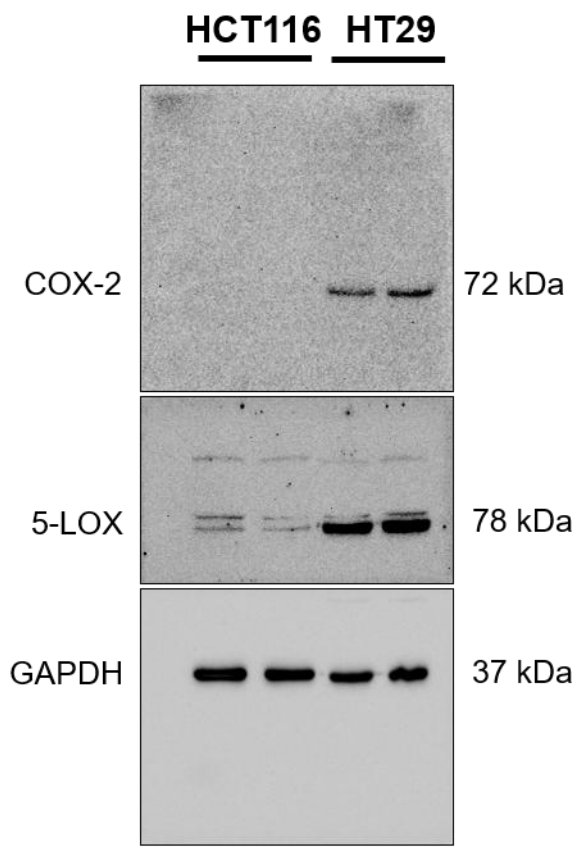

Fig. 1 E

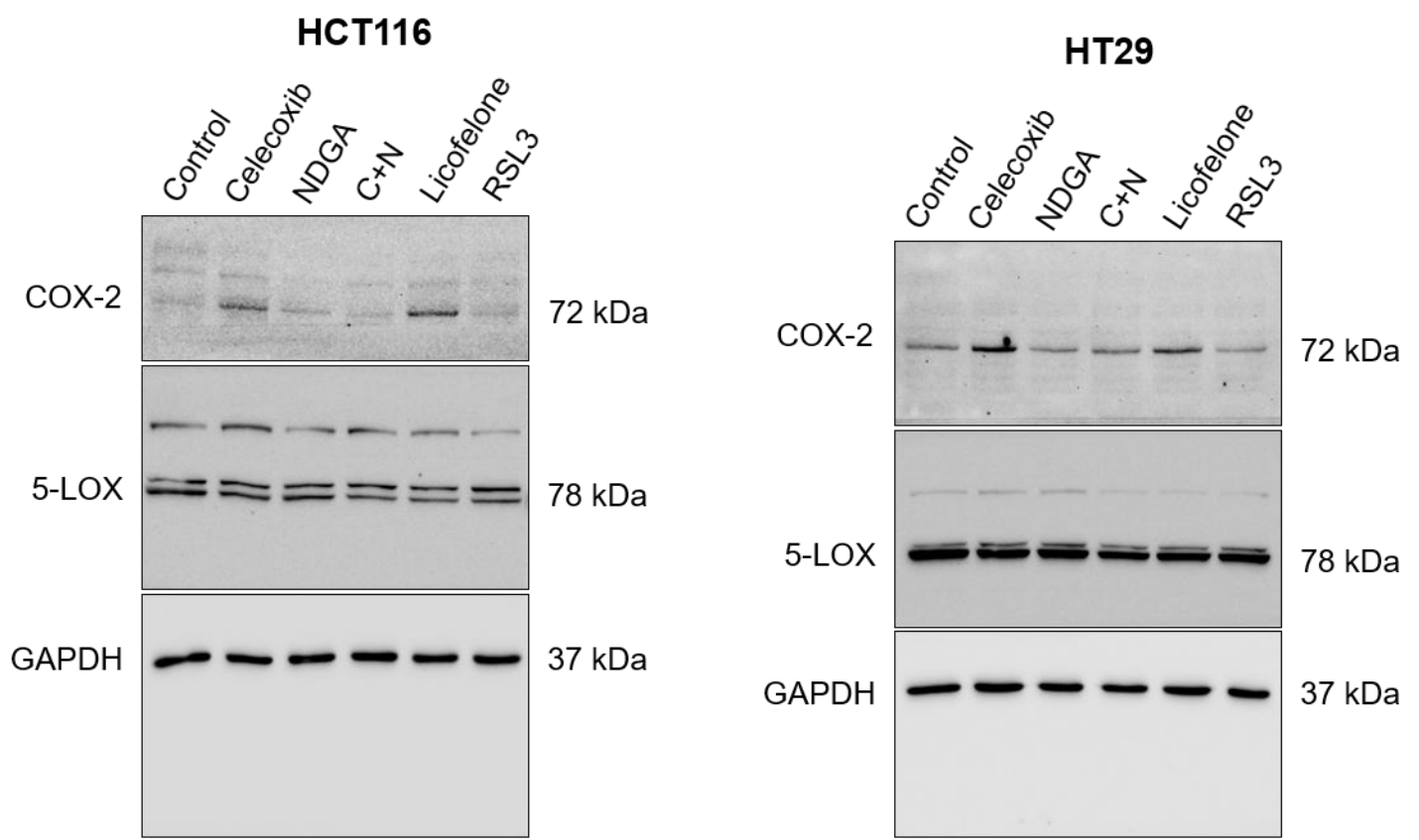

**Fig. 2 B**

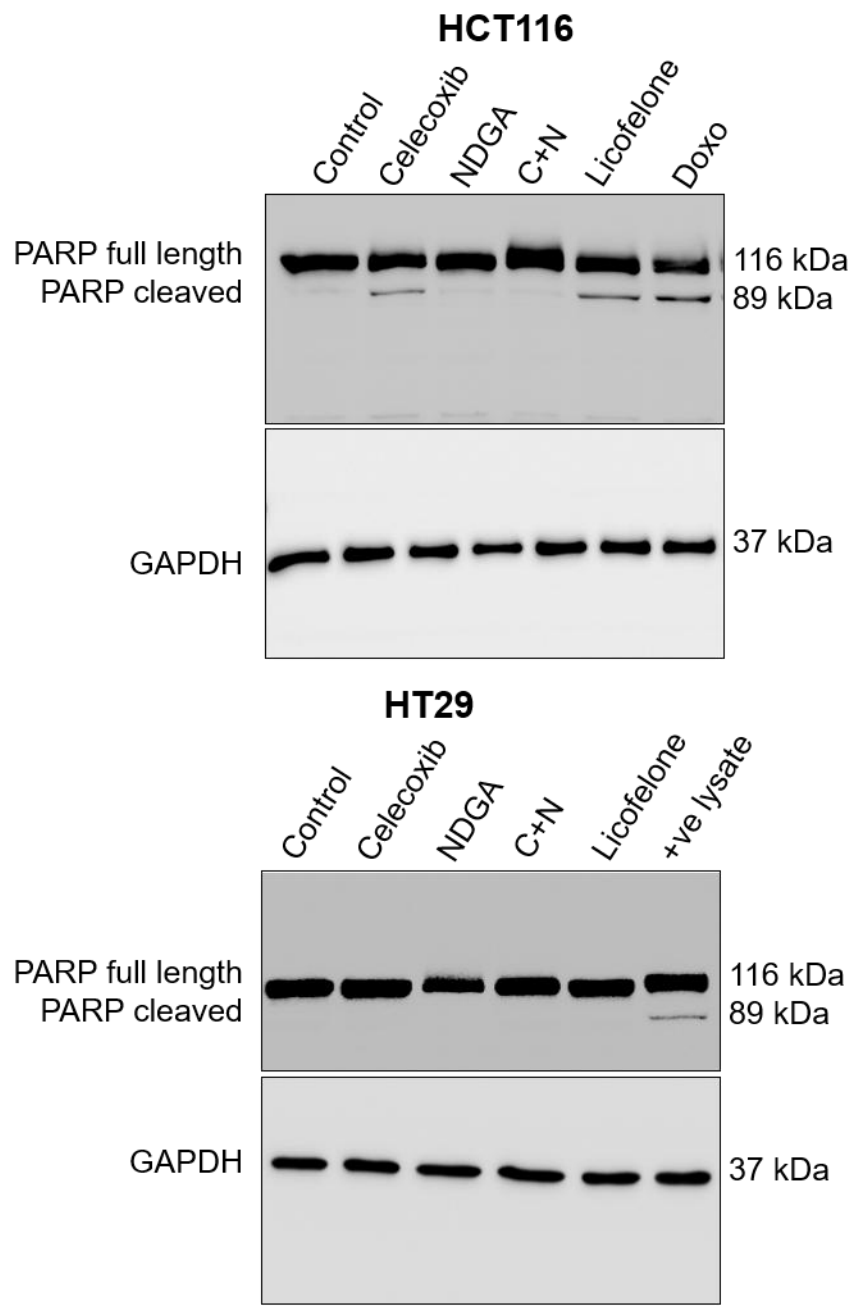

**Fig. 6**

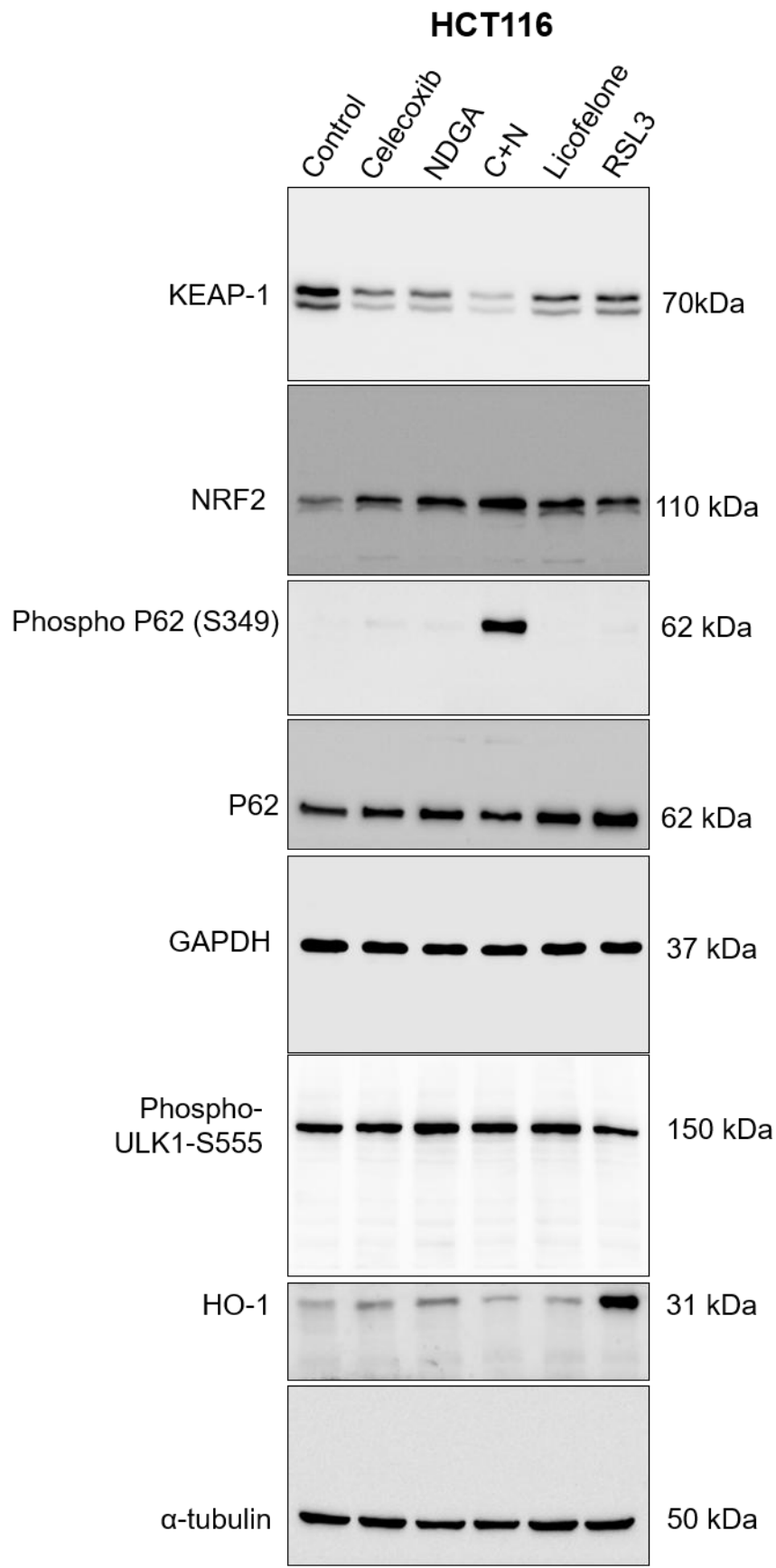

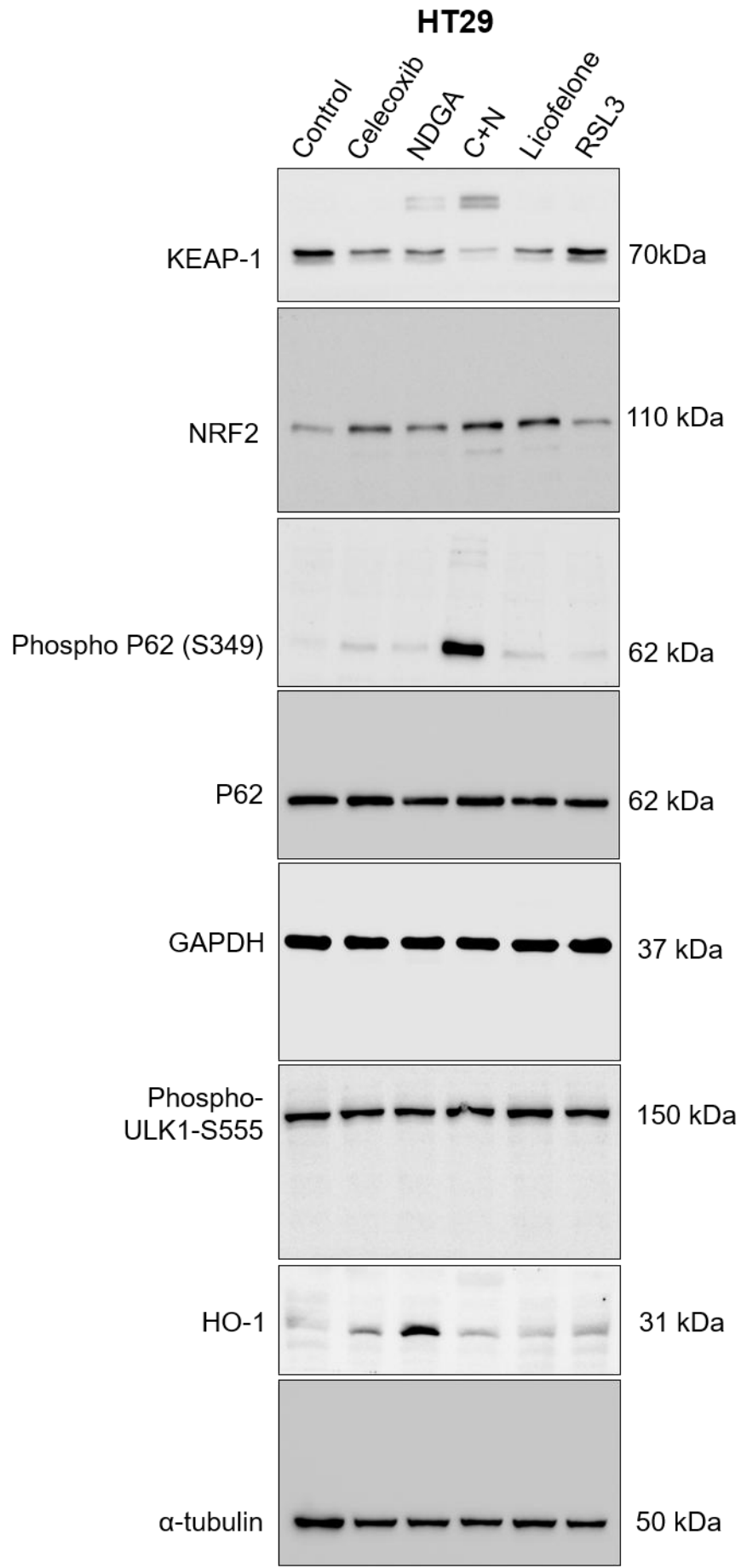
