## Supplemental data 2 for "Combined celecoxib and NDGA induces a non-canonical, autophagy-independent redox cell death via p62/KEAP1 in colorectal cancer"

**Fig. S1**

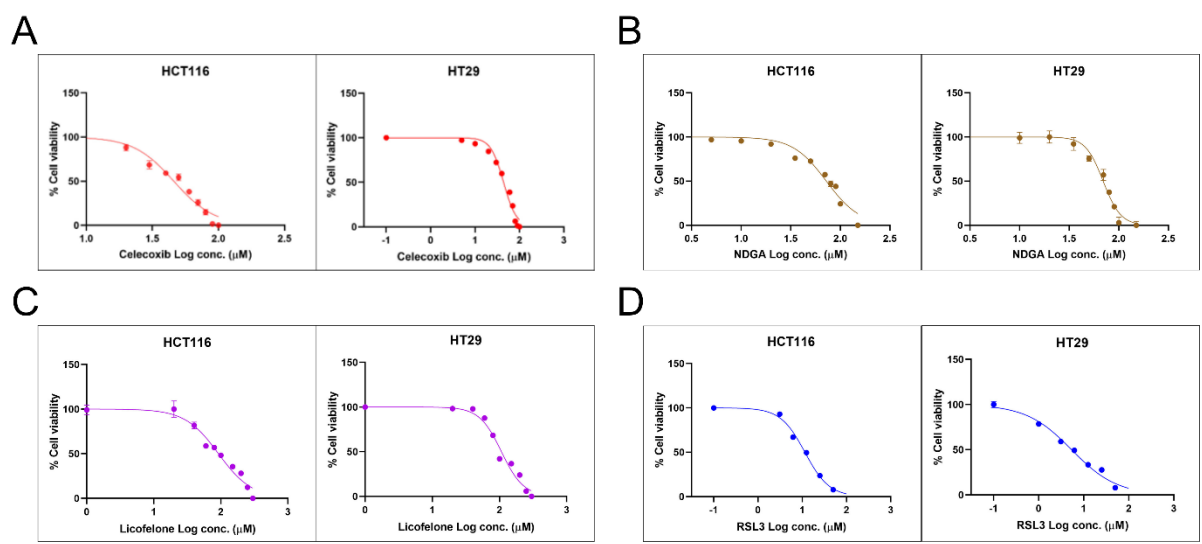

**Figure S1. Dose-response curves of (A) NDGA, (B) Celecoxib, (C) Licofelone, and (D) RSL3 in HCT116 and HT29 cell lines.** Cells were treated with increasing concentrations of indicated compounds for 24 hrs. Data were presented as mean ± SEM from at least three independent experiments performed in triplicate. Sigmoidal dose-response curves were plotted using a four-parameter logistic (variable slope) model. Calculated IC<sub>50</sub> values are summarized in Supplementary figure Table 1.

**Table S1.**

| Inhibitors | IC<sub>50</sub> (μM) ± SEM<br>HCT116 | IC<sub>50</sub> (μM) ± SEM<br>HT29 |
| --- | --- | --- |
| Celecoxib | 45.94 ± 1.40 | 43.76 ± 1.29 |
| NDGA | 72.23 ± 2.11 | 69.51 ± 1.51 |
| Licofelone | 92.16 ± 3.94 | 106.8 ± 3.38 |
| RSL3 | 11.49 ± 0.37 | 5.38 ± 0.37 |

**Fig. S2**

**A**

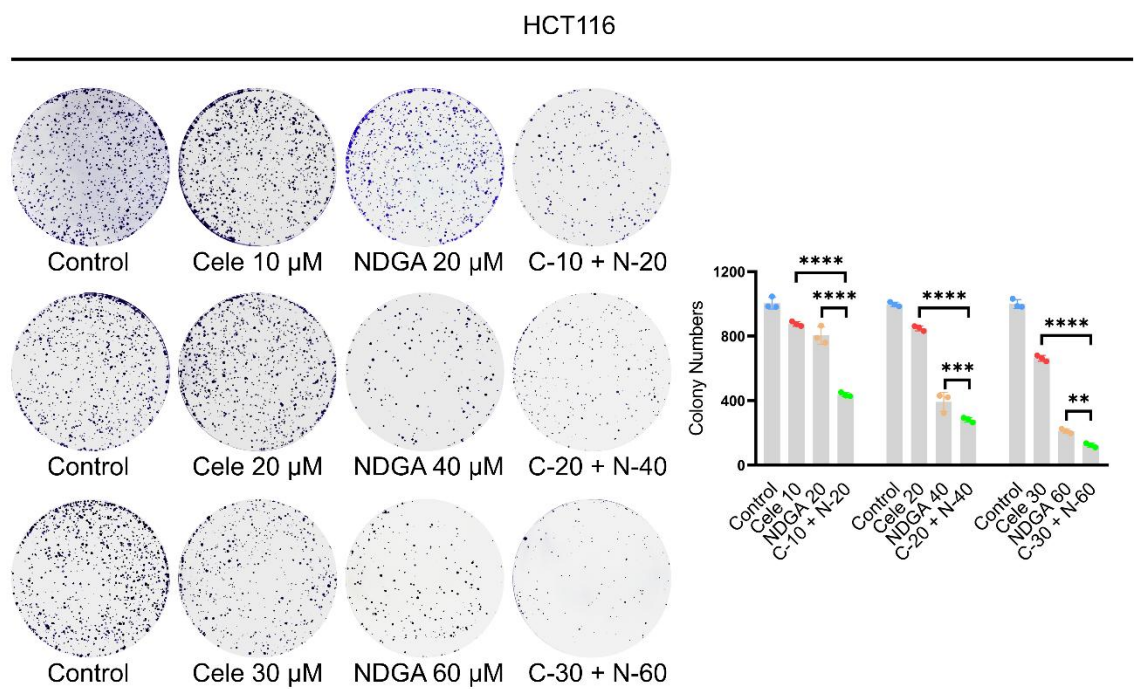

**B**

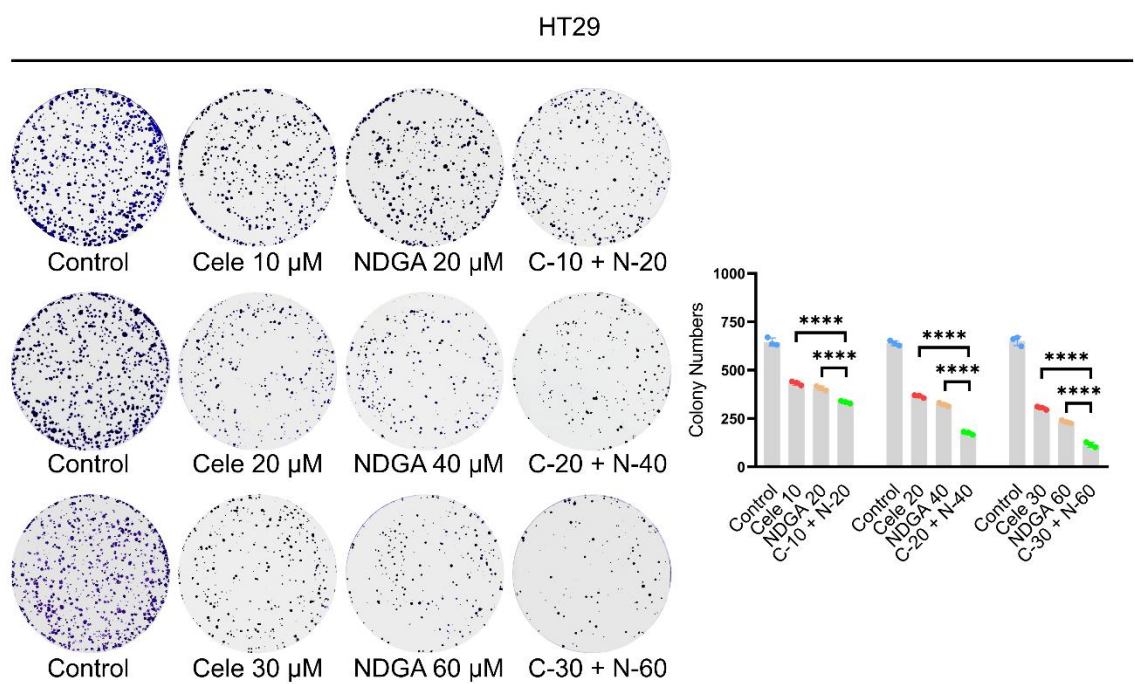

**Figure S2. Representative figures of clonogenic assay in varying concentrations of celecoxib, NDGA, and their combination (A) HCT116** Colony formation assay with the indicated treatments was performed (left panel), and a histogram of colony numbers is shown (right panel). Data are shown as mean ± SD (n=3) (\*\*p < 0.005; \*\*\*p < 0.0005; \*\*\*\*p < 0.0001 versus celecoxib alone or NDGA alone). **(B).** HT29 Colony formation assay with the indicated treatments was performed (left panel), and a histogram of colony numbers is shown (right

panel). Data are shown as mean ± SD (n=3) (\*\*\*\*p <0.0001 versus celecoxib alone or NDGA alone).
